# An Interpretable 3D Bag-Of-Visual-Words Pipeline for Volumetric Microscopy Classification

**DOI:** 10.64898/2026.04.21.719969

**Authors:** Anna E. Pittman, Kirby R. Campbell, Christophe Laumonnerie, David J. Solecki

**Affiliations:** Neuronal Cell Biology Division, Department of Developmental Neurobiology, St. Jude Children’s Research Hospital, 262 Danny Thomas Place, Memphis, TN 38104, USA

## Abstract

Fluorescence microscopy increasingly produces complex volumetric datasets whose biologically meaningful differences are difficult to capture with hand-crafted measurements, especially when signal is distributed across three-dimensional space. Here, we present an interpretable 3D Bag-of-Visual-Words (BoVW) pipeline for classification and analysis of volumetric microscopy data. The framework detects multiscale local keypoints, computes rotationally robust 3D gradient-based descriptors, and aggregates them into image-level visual-word representations. These features are then used for low-dimensional visualization and logistic regression classification, while model weights are mapped back to the original volumes to generate attention maps that localize discriminative structures. We applied the pipeline to two cerebellar granule neuron datasets spanning both ideal and non-ideal imaging conditions. In a near-isotropic lattice light-sheet dataset of chromatin organization, the method separated control and NIPBL loss-of-function nuclei and supported accurate classification, with strongest performance in the facultative heterochromatin and H3.3 channels. Attention mapping and downstream connected-component and Haralick analyses revealed that loss-of-function nuclei contained more fragmented high-attention regions and smoother, more homogeneous chromatin-associated textures than controls. We then evaluated the same framework on an anisotropic confocal timelapse dataset of receptor clustering in dense neuronal cultures, where single-cell segmentation was impractical. Despite these challenges, the representation captured the expected ligand-driven clustering response and resolved subtler differences associated with a polarity protein overexpression. Together, these results establish a simple, interpretable, and broadly applicable framework for extracting biologically meaningful structure from volumetric microscopy datasets while preserving native 3D context.

## 1. Introduction

Fluorescence microscopy has transformed the study of biological systems, enabling direct visualization of subcellular structures and dynamics. However, the increasingly sophisticated imaging strategies that drive biological discovery also produce datasets that are large, heterogeneous, and difficult to analyze^1^. 2D datasets, such as whole-slide histology images, can be challenging due to their large individual sizes, while high-throughput experiments generate tens of thousands of smaller fields of view that rapidly overwhelm manual and semi-manual workflows. These challenges are further compounded in volumetric datasets and in modalities that probe intrinsically three-dimensional structures, for which open source, general-purpose analysis tools are comparatively scarce.

A common approach for analyzing 3D images is to treat them as stacks of 2D slices. This is often done to reduce computational demands, simplify visualization, and mitigate anisotropy artifacts that are common along the z-dimension in microscopy data. For example, open-source tools such as Cellpose typically segment volumetric data by running 2D models on individual planes and then reconstructing 3D objects from the slice-wise predictions^2^. While this approach has been successful and efficient for segmentation, it is less suitable for tasks that depend on genuine 3D structure, such as feature extraction and classification of nuanced fluorescence signal. Down sampling or flattening the data to 2D can discard important information about spatial relationships and topology. There is a need for methods that operate directly on volumetric images, generating rich, interpretable feature representations without reducing the data to lower dimensions prematurely.

Interpreting volumetric fluorescence data is inherently difficult for human observers. Meaningful visualization typically requires reducing 3D information to 2D slices or projections, but this collapse discards spatial context and can obscure relationships that exist only in three dimensions. As a result, hand-crafted analyses often default to coarse summary metrics (e.g., overall size or shape anisotropy) that capture only a small fraction of the available signal. There is therefore a need for methods that preserve native dimensionality while providing interpretable summaries that an expert can use to refine—or challenge—existing biological interpretations.

Here, we present a 3D analysis framework based on a Bag-of-Visual-Words (BoVW) pipeline with a custom 3D feature extractor. Local 3D fluorescence signal structure is summarized using rotationally robust descriptors, and these descriptors are aggregated into an image-level BoVW representation. The resulting feature vectors are used to train a logistic regression classifier that distinguishes between relevant conditions and, via its learned weights, highlights intensity patterns and spatial motifs that are most informative for classification.

We focus on cerebellar granule neurons (CGNs), an ideal system for method development because their maturation is tightly regulated and accompanied by stereotyped, well-characterized morphological transitions. In the developing cerebellum, CGN progenitors reside in the external granule layer (EGL) where they undergo clonal expansion, then exit the cell cycle, differentiate, and migrate inward to form the mature granule layer^3^. This progenitor-to-differentiated transition is marked by two hallmark events central to our study: extensive chromatin reorganization during differentiation^4^, and the proper direction of migratory behavior required for timely exit from the EGL^5^. Disrupting either process has major consequences—prolonged proliferation in the EGL is associated with medulloblastoma^6^, while premature cell-cycle exit can reduce CGN number and affect proper lamination of the cerebellum, which has been linked to neurodevelopmental disorders^7,8^.

These behaviors can be read out with interpretable fluorescence markers, including chromatin markers that report nuclear organization and cell-surface guidance receptors that report migration-related signaling necessary for proper migratory direction and progression out of the EGL. Our 3D nuclear dataset targets chromatin organization using four different chromatin markers imaged with lattice light-sheet microscopy, whose near-isotropic resolution, a rarity in the microscopy world, makes the volumetric structure well suited for true 3D analysis. The variety of chromatin markers allows us to investigate which markers contain condition-specific changes and gives us a deep understanding of the underlying biology.

Our 3D timelapse dataset captures receptor clustering dynamics before and after ligand addition, in both control cells and cells with perturbation of a polarity pathway. This design provides two complementary scales of phenotypic change: a strong, visually striking ligand-driven clustering response and a subtler modulation associated with the genetic perturbation. The dataset also reflects common constraints of dense neuronal cultures—extensive overlap and intermingling processes make single-cell segmentation and cropping impractical. Additionally, it also reflects common microscopy constraints: confocal acquisition yields pronounced axial anisotropy and sampling steps chosen for speed lead to sparse axial sampling, compromising the resolution in the axial dimension. Together, these factors create a stringent test of algorithmic robustness under realistic, non-ideal imaging conditions that deviate from clean, isotropic volumes.

A central design principle throughout this work is minimal complexity. We aim to build the simplest pipeline that adequately addresses a given dataset, adding additional layers of complexity only when the data requires them. The BoVW backbone was chosen in order to pursue the most transparent simple pipeline; we avoid layering on neural networks and opt instead for a simple logistic regression model to dig deeper into the data.

The full 3D image analysis pipeline—rotationally robust local descriptors, BoVW aggregation, and a linear classifier with mappable weights—enables condition separation while retaining 3D context and interpretability in both the ideal (isotropic single cell crops) and more realistic (many overlapping cells and anisotropic acquisition) use cases. Ultimately, it’s a broadly applicable framework designed to generalize across a variety of fluorescent datasets.

## 2. Related Works

Bag-of-Visual-Words (BoVW), inspired by bag-of-words models in text^9^, represents an image by quantizing local intensity descriptors into a dictionary of “visual words” and summarizing each image as a histogram of those words^10^. These histograms form compact feature vectors that support tasks such as image retrieval, clustering, and classification. They are often visualized using dimensionality reduction methods such as principal component analysis (PCA) or uniform manifold approximation and projection (UMAP). Our method adopts the BoVW paradigm for its interpretability and modularity, allowing us to design the local descriptor and sampling strategy to properly reflect genuinely 3D structures from biological fluorescent volumetric imaging whereas standard BoVW pipelines often rely on 2D inputs, or 3D images that have been broken down into 2D slices.

BoVW has been applied to biomedical imaging in multiple contexts, particularly in 2D pathology where local tissue morphologies recur across large cohorts. For instance, Cruz-Roa et al. use BoVW-style representations for histology analysis, demonstrating that dictionaries of local patterns can capture meaningful visual content in HCE imagery^11^. Related unsupervised “visual phenotype” approaches use dictionaries to discover disease-associated morphologies and link them to clinical or molecular endpoints. Powell et al. construct image-derived visual words from TCGA glioma slides to predict survival and associate predictive phenotypes with signaling activity, illustrating how dictionary-based representations can bridge image patterns to biological hypotheses^12^. Similarly, Lee et al. propose an unsupervised bag-of-words framework on renal biopsy whole-slide images to predict functional outcomes^13^, while noting that missing spatial information can contribute to misclassifications—highlighting a recurring tradeoff between compactness and spatial specificity. In contrast to these primarily 2D, tissue-scale settings, our focus is cellular or subcellular organization in volumetric fluorescence microscopy, where discriminative cues arise from 3D structures within the images.

BoVW has also been applied to CT volumetric data; Feulner et al. apply a bag-of-words representation to CT volumes for body-part estimation, demonstrating that local descriptors can compactly represent 3D scans^14^. However, that analysis broke down the 3D image into 2D slices before feature extraction. While that was sufficient for the analysis of CT volumes, volumetric fluorescence microscopy generates complex 3D structures that must be analyzed in their native dimensionality. Our approach addresses these microscopy-specific constraints by extracting local 3D descriptors within each image and aggregating them into an image-level BoVW vector used for classification and interpretation.

A key challenge in adapting BoVW to fluorescence volumes is handling arbitrary orientation of subcellular structures. Prior work on rotation robustness typically relies on canonical orientation assignment, orientation pooling, or expressing gradients in a relative reference frame. Our descriptor follows the latter strategy: we adapt a rotation-invariant histogram of oriented gradients (HOG) variant that replaces absolute gradient orientation with locally relative measurements^15^ and extend the formulation to 3D microscopy volumes (Methods).

Taken together, prior work shows that dictionary-based representations can summarize complex imagery and support downstream prediction, including in biomedical and even volumetric settings. At the same time, standard BoVW pipelines rely on 2D inputs either from 2D images or 2D slices of 3D volumes. This motivates our framework: a BoVW pipeline designed for volumetric fluorescence signal, using rotationally robust 3D local descriptors and a linear classifier whose weights can be mapped back to informative intensity patterns and spatial motifs.

## 3. Datasets and Problem Setup

### Problem definition

Traditional analyses rely on an expert observer to recognize global trends and then design concrete measurements that capture those trends. In some cases, this is straightforward—for example, measuring cell velocity for a motility phenotype. However, as the underlying signal becomes more nuanced with less obvious trends, it is increasingly difficult to identify a single, hand-crafted quantity that adequately summarizes the phenotype. This is especially true for structural properties such as chromatin organization, where the phenotype might be visually apparent but not easily reducible to a small set of intuitive descriptors.

A further challenge arises when the signal is inherently three-dimensional. For a human observer, it is extremely difficult to meaningfully interpret volumetric structure without collapsing the data into 2D slices or projections. Although these projections can be informative—and may even be sufficient in some settings—they capture only a fraction of the information present in the volume, leaving much of the signal unexamined.

Computational methods do not face the same constraint. Models can operate directly on the data in its native dimensionality, enabling them to interrogate the full 3D structure rather than a compressed projection. This makes it possible to leverage volumetric information that is difficult to access by eye and to reveal trends that might otherwise remain hidden.

Our goal is to complement careful expert inspection with a flexible, data-driven representation that can capture phenotypic differences without requiring an *a priori* choice of a relevant measurement. By operating directly on volumetric data in its native dimensionality, this approach preserves 3D spatial context and enables the analysis to leverage structural information that is difficult to articulate or quantify by hand. To this end, we build on a Bag of Visual Words (BoVW) backbone and develop a pipeline that *(i)* isolates image structures that differ between conditions and *(ii)* provides interpretable summaries of those structures.

### Dataset 1: 3D chromatin and nuclear architecture

We analyzed a live-cell volumetric dataset designed to probe nuclear and chromatin organization during CGN differentiation. Samples consist of multi-channel 3D images providing complementary readouts of nuclear architecture and chromatin state (including chromatin-associated markers and a DNA label). Two conditions are analyzed, control neurons and Nipped-B-like protein (NIPBL) loss of function (LOF) neurons. NIPBL is part of the cohesin complex, responsible for loading cohesin onto the DNA. The loss of function of that protein disrupts chromatin structure and leads to an increase of nuclear volume. The objective is to distinguish these conditions based on subtle, spatially distributed changes in 3D nuclear organization. This dataset directly targets one of the central biological features of CGN maturation—chromatin reorganization—while preserving the full volumetric context needed to capture higher-order structure.

### Dataset 2: 3D signaling and membrane receptor clustering dataset

We evaluated the robustness of the pipeline on a three-dimensional confocal timelapse dataset of cerebellar granule neurons that reports changes in cell-surface signaling organization. In contrast to the nuclear dataset, these volumes are strongly anisotropic with comparatively coarse axial sampling due to the nature of confocal acquisition and above-Nyquist sampling in the lateral dimensions chosen to increase sampling speeds. Moreover, the high neuronal density precluded reliable single-cell segmentation and made it impractical to crop volumes to individual cells. As a result, each field of view unavoidably contains substantial biological heterogeneity (e.g., mixtures of different plasmid expression levels or differentiation status), providing a stringent test of whether the method can extract condition-associated structure without cell-level normalization.

The images capture the spatial distribution of a guidance receptor at the plasma membrane before and after ligand addition, a manipulation known to induce receptor clustering. Samples were labeled by experimental condition (control versus a perturbation). This dataset is an informative use case because the phenotype is biologically meaningful and readily recognizable by eye, yet it is expressed through distributed, local changes in receptor organization rather than a single obvious scalar measurement.

### Key challenges and analysis objectives

Across both datasets, the core difficulty is that the discriminative signal is often structural, distributed, and not easily summarized by a small set of hand-crafted descriptors. The high dimensionality of the data makes manual feature design and interpretation particularly challenging. This is compounded in the timelapse dataset featuring high density of overlapping cells, anisotropy inherent in confocal microscopy, and acquisition parameters optimized for speed leading to lower resolution in the z-dimension.

To address these constraints, we used a Bag-of-Visual-Words (BoVW) backbone to form image-level summaries from collections of local patterns. We utilized this framework combined with a custom 3D descriptor and trained a lightweight linear model on image–label pairs. Importantly, the learned model weights are used to generate attention maps that highlight which local intensity patterns within each image or volume contribute most strongly to the classification, supporting interpretation alongside discrimination.

## 4. Method

### 4.1 Overview

We developed a 3D BoVW pipeline (Fig. 2) to summarize volumetric fluorescence patterns with an interpretable, image-level representation. Starting from either whole FOV images or single-cell 3D crops with corresponding masks, we detect multi-scale local keypoints, compute rotationally robust 3D gradient-based descriptors, and encode these descriptors using a learned visual dictionary (whose size is set to 200 words). Descriptor encodings are pooled within each image to produce a 200-dimensional BoVW vector per image, which is used for visualization and for supervised classification of experimental condition. To interpret model decisions, we back-project the linear classifier’s contributions from visual words to the underlying patches, generating volumetric attention maps that localize discriminative nuclear structures. We further quantify the spatial organization and texture of high-attention regions using 3D connected-component (“blob”) analysis and Haralick texture features.

**Figure 1.**
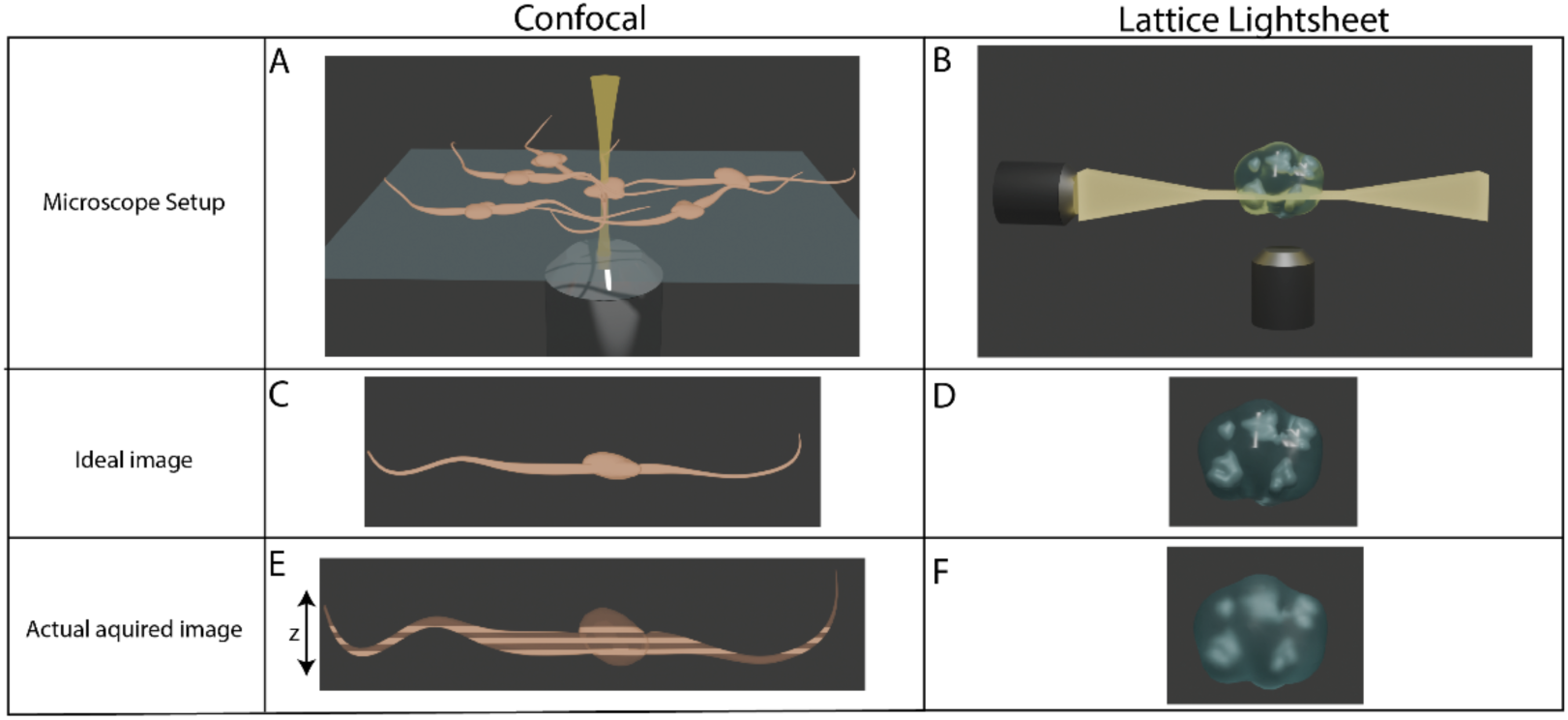
Dataset Acquisitions. Cartoon illustrations show microscope setups and outputs. Spinning disk confocal microscopes image fast but su<er from anisotropy in the z-dimension leading to aberrant elongation [A, C, E]. Large step sizes in the z-dimension lower axial resolution [see striping in E]. Lattice light-sheet illuminates the sample with a thin sheet, optical setup leads to near isotropic capture of the sample [B, D, F].

**Figure 2.**
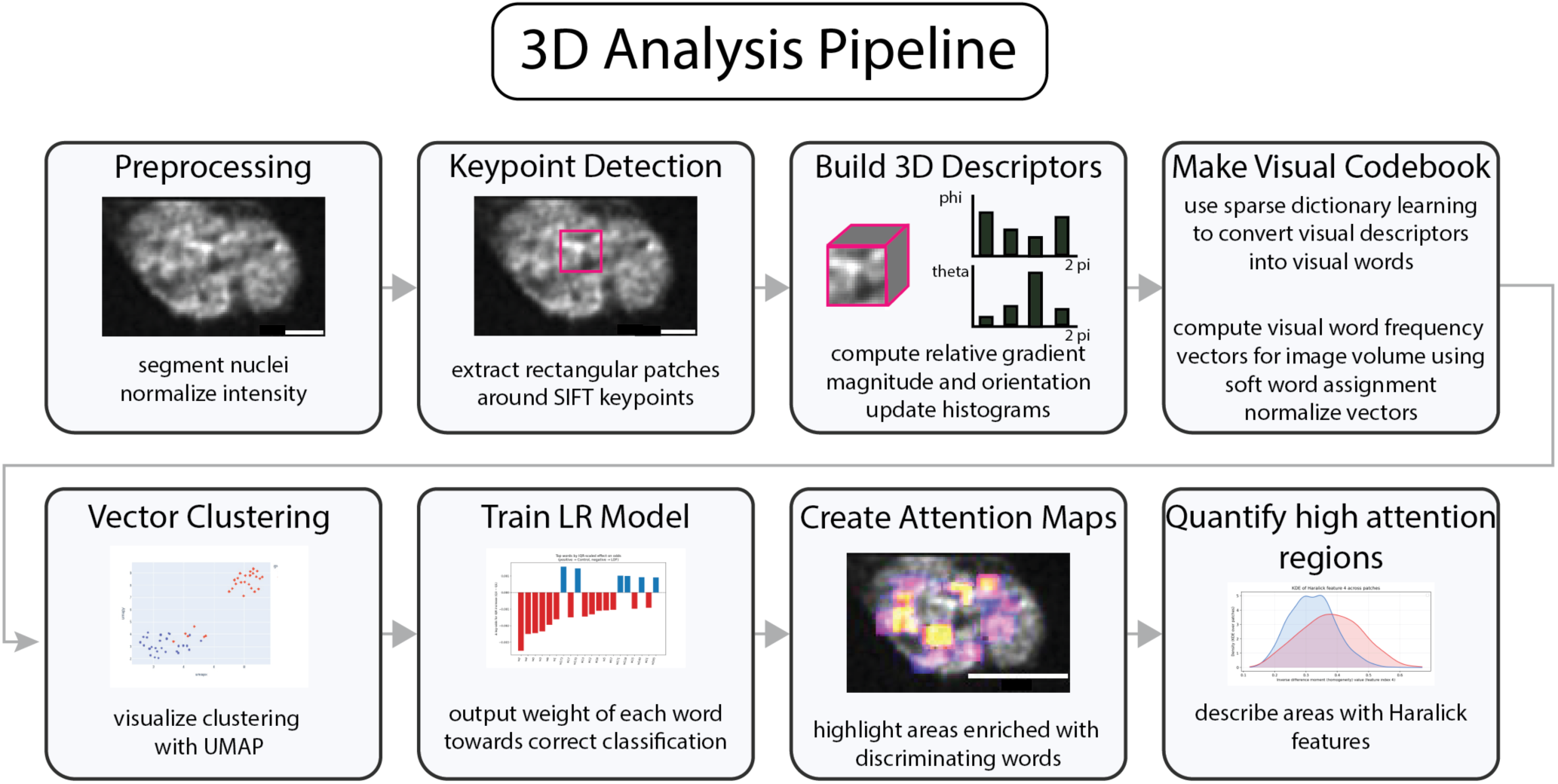
3D analysis pipeline. Volumetric images are processed through the pipeline starting with the preprocessing step. Scale bar = 2 µm.

### 4.2 3D Bag of Visual Words Representation

Cultured primary CGNs were nucleofected with indicated imaging probes or biosensor as previously described^16–19^ and volumetric image stacks were acquired either using the lattice light-sheet microscopy or spinning disk confocal. Intensities were normalized per nucleus by clipping to the 5th–95th percentiles and rescaling to the [0,1] range. Other normalization protocols were assessed and did not impact the final conclusions (SI Figure 7). For the LLSM images, nuclear segmentation masks were generated using Cellpose SAM^20^, single-cell crops were generated using the masks, and all subsequent feature extraction was restricted to voxels within the nuclear mask. For the timelapse data, entire FOV images were used with no corresponding 3D cell masks.

Local keypoints were detected using a 3D implementation of the Scale-Invariant Feature Transform (SIFT) algorithm^21^, which starts by computing a difference-of-Gaussians (DoG) scale space. For each image, the normalized volume was convolved with 3D Gaussian kernels at multiple scales σ ∈ {1.0,1.6,2.2,3.0}. Differences between successive Gaussian-blurred volumes formed a set of DoG volumes that approximate the Laplacian-of-Gaussian. This procedure was repeated over three octaves, each obtained by down sampling the volume by a factor of two, yielding a multi-scale 3D DoG pyramid.

Within each DoG volume, candidate keypoints were voxels that were local extrema in a 3×3×3 neighborhood whose DoG response exceeded a user-defined contrast threshold in absolute value. We retained extrema corresponding to bright-blob structures (DoG minima) and dark-blob structures (DoG maxima). Candidate coordinates were then mapped back to the original image resolution according to their octave (multiplication by 2^octave^). To suppress edge-like responses, we applied a 3D analogue of the SIFT edge filter: for each candidate, the 3×3×3 DoG neighborhood Hessian was estimated, its eigenvalues were computed, and points with a large anisotropy ratio (largest/smallest eigenvalue greater than the user-defined threshold) were discarded. Each retained keypoint was thus characterized by its 3D location in the original volume and by a discrete scale index (octave and σ-level).

For each keypoint, we computed a 3D histogram-of-oriented-gradients (HOG) descriptor, adapted to Python from the Matlab implementation of Tzimiropoulos et al^22^. The 3D intensity gradient was obtained by finite differences along the x, y, and z axes of the normalized image. Within a cubic patch centered at the keypoint, with edge length proportional to the SIFT scale (5σ), each gradient vector was converted to spherical coordinates, and its magnitude was accumulated into histogram bins defined over the polar (θ) and azimuthal (φ) angles. Voxels were included only if a sufficient fraction of the patch lay inside the nuclear mask, ensuring that descriptors captured nuclear rather than background structure.

Standard HOG descriptors are not rotationally invariant: rotating an image changes the absolute orientation of gradients and thus the resulting histograms. To mitigate this, we adapted the rotationally invariant HOG formulation of Cheon et al. from 2D to 3D^15^. Instead of using absolute gradient orientations, we expressed each voxel’s gradient magnitude and orientation relative to those of its 3D neighborhood (26-connected neighbors). Relative orientation differences (in θ and φ) and relative magnitude ratios were accumulated into the orientation histograms. Because these quantities are defined with respect to local neighbors rather than the global coordinate frame, the resulting descriptors are approximately invariant to rigid rotations of the nucleus.

Descriptors from all images were pooled to learn a 3D visual codebook. A 200-word codebook was constructed using sparse dictionary learning (`MiniBatchDictionaryLearning` from scikit-learn). The size of the codebook was empirically determined to best capture the full range of data. This procedure learns a set of basis vectors (“visual words”) such that each descriptor can be approximated as a sparse linear combination of these words. We used soft assignment: each keypoint descriptor was encoded as a vector of coefficients over the entire dictionary, rather than being assigned to a single word.

For each image, patch-level coefficient vectors were summed to obtain an image-level representation, yielding a 200-dimensional vector of word “frequencies” or usage weights. To emphasize informative words and down-weight ubiquitous ones, these image-level vectors were transformed using a term-frequency–inverse-document-frequency^23^ (TF–IDF) scheme and then L2-normalized. The L2 normalization also reduced sensitivity to differences in the number of keypoints per cell (e.g., due to cell size). These normalized 200-dimensional vectors constituted the primary representation used for all downstream analyses.

For visualization, we embedded the normalized cell-level vectors into a low-dimensional space using UMAP. Unless otherwise specified, standard UMAP parameters were chosen to qualitatively preserve both local and global neighborhood structure.

For the nuclear dataset, in order to partially control for size-related heterogeneity, the visual codebook was learned on the set of descriptors from the entire dataset, but downstream analyses were conducted on size-stratified subsets of the data. Specifically, we focused on larger LOF nuclei (> median nuclear volume) and smaller control nuclei (< median nuclear volume), with volumes derived from the 3D nuclear segmentations (for analysis on non-stratified datasets see the Supplementary Information). NIPBL knockdown led to increased nuclear volume; enriching for larger nuclei therefore enriched for cells most affected by the perturbation. For the timelapse dataset the dictionary was learned and applied to the entire dataset.

### 4.3 Logistic Regression Classifier

We trained a logistic regression classifier on the normalized BoVW vectors to quantify the extent to which the representation captured the phenotype. The input features were the 200-dimensional normalized frequency vectors, and the target labels were the known experimental conditions (*i.e.* control vs. LOF). Logistic regression was used with standard L2 regularization. For the nuclear images a two-class model was trained (control vs LOF) and for the membrane receptor images a four-class model was trained to account for control and overexpression pre and post addition of ligand. Model performance for the two-class model was summarized by the area under the receiver-operating characteristic curve (AUC-ROC), computed from the predicted probabilities across cells.

The fitted model provided a coefficient for each visual word, indicating the degree and direction with which that word contributed to predicting the correct class. Words that pushed towards the correct class were given positive weights, words which pushed towards the incorrect class (confounding information) were given negative weights.

### 4.4 Attention Map Construction

To localize discriminative information within each image, we decomposed the linear classifier’s image-level decision value into patch-level contributions. For each image, we first reconstructed the TF–IDF and L2-normalized visual word vector used during logistic regression training and computed the contribution of each word to the decision value as *w*_*k*_*x*_*k*_, where *w*_*k*_is the learned logistic regression coefficient and *x*_*k*_is the normalized TF–IDF weight of word *k*. We then distributed each word’s contribution back to individual patches in proportion to the absolute usage of that word in each patch, such that patches containing more of a given word received a larger share of that word’s contribution. Summing these contributions over all words yielded a scalar attention score for each patch. By construction, the sum of patch scores equals the image-level classifier score (up to numerical precision), providing a faithful spatial decomposition of the linear decision function.

For intuition, consider a visual word that contributes 10 arbitrary units to the classifier score for a given nucleus. If this word is distributed equally across 10 patches (each patch carrying the same absolute usage of that word), then each of those patches receives 1 unit of that contribution, and their patch-level contributions sum to the original 10 units.

Patch-level attention scores were then mapped back onto the 3D image volume using the original patch locations, producing a volumetric attention map for each nucleus. Attention maps were rescaled on a per-nucleus basis (e.g., min–max normalization) to facilitate visualization and comparison across images.

### 4.5 Nuclear Blob and Texture Analysis

To quantify the spatial organization of high-attention regions, the attention maps were thresholded to define a binary mask of “high-attention” voxels. A single global threshold was chosen empirically based on the distribution of attention scores, and this threshold was applied uniformly across nuclei.

Within each binary attention map, connected-component labeling in 3D was used to identify isolated high-attention “blobs.” For each nucleus, we recorded (*i*) the number of blobs and (*ii*) the volume (voxel count) of each blob. These measurements were compared between conditions to assess differences in the fragmentation and spatial extent of the most informative regions.

To characterize the local intensity patterns within high-attention regions, we computed Haralick texture features on 3D patches. For each image, voxel intensities within the nuclear mask were clipped to the 1st–99th percentiles, normalized to [0,1], and quantized to 32 gray levels, with background voxels set to zero. High-attention patches (as defined by the attention threshold) were extracted from these quantized volumes, and Haralick features were computed using the Mahotas Python package (`mh.features.haralick, distance = 1, ignore_zeros = True`). This function constructs gray-level co-occurrence statistics over multiple directions and, with `return_mean = True`, returns the standard set of 13 Haralick features averaged over directions for each patch.

Each high-attention patch was summarized by a 13-dimensional Haralick feature vector. We analyzed these features at two levels. First, for each nucleus we computed the mean value of each Haralick feature across its high-attention patches, yielding one 13-dimensional summary vector per image. To examine the full distribution of texture values across high-attention regions, we pooled patch-level feature values across all images within a condition and estimated kernel density distributions (KDEs) for individual Haralick features.

## 5. Experiments and Results

### 5.1 Experimental Setup

We evaluated the proposed BoVW framework on two microscopy datasets (*i*) a 3D chromatin architecture dataset used to test whether the 3D BoVW representation supports both condition discrimination and localization of discriminative structure within nuclei and (*ii*) a 3D timelapse receptor clustering dataset, which tests robustness on a realistic acquisition where images are anisotropic and cannot be reduced to single-cell crops.

#### 3D chromatin dataset (segmented single-nucleus volumes)

In the chromatin dataset, each sample corresponded to a segmented and cropped nucleus, enabling direct cell-level comparison across conditions (control vs. NIPBL knockdown). The primary analysis focused on a custom designed H3K27me3 H2AK119ub biosensor for facultative heterochromatin (manuscript in preparation), and the same workflow was also applied to additional markers (Hoechst, H3.3, and CTCF). Each probe was chosen to focus on different types of chromatin. The facultative heterochromatin probe accumulates in facultative heterochromatin, which is heterochromatin whose accessibility can be developmentally regulated. H3.3 is associated with transcriptionally active chromatin, CTCF is enriched at topologically associated domain boundaries, and Hoechst binds the minor groove preferentially in AT-rich regions.

Because biological variability in knockdown extent introduced heterogeneity, we performed downstream analyses on size-stratified subsets (larger LOF nuclei and smaller control nuclei, split at the median nuclear volume) to enrich for cells most affected by the perturbation while reducing overlap driven by size-related variability.

To quantify the spatial organization of discriminative regions, attention maps were thresholded to define high-attention voxels, followed by 3D connected-component analysis to measure the number and size distribution of high-attention “blobs” per nucleus. To further characterize local structure within these regions, we computed 3D Haralick texture features on high-attention patches and compared feature distributions between conditions.

#### 3D timelapse receptor dataset (frame-wise field-of-view volumes)

For the receptor clustering timelapse dataset, sequences were split into individual timepoints, yielding a series of 3D frames treated as independent samples for representation learning and classification. Unlike the chromatin dataset, individual cell segmentation and cropping were infeasible due to high cell density and overlap, and the confocal acquisition produced pronounced axial anisotropy, creating a challenging “non-ideal” test case. Labels captured both a strong ligand-driven clustering response (pre-vs post-ligand) and a subtler modulation associated with pathway perturbation.

### 5.2 3D Chromatin Architecture Results

Previous work in the lab attempted to distinguish control and NIPBL loss-of-function (LOF) conditions using hand-crafted features derived from expert visual inspection. Here, we sought to identify similar or additional trends using an unsupervised Bag of Visual Words approach adapted to 3D data. The normalized frequency vectors exhibited clustering behavior, (Figure 3B), that separated control and NIPBL LOF cells. Some overlap between conditions as seen, consistent with biological variability in the degree of knockdown, differences in differentiation status, and other sources of heterogeneity. However, when size-based stratification was applied, the separation between control and LOF clusters became more pronounced (SI Figure 4).

**Figure 3.**
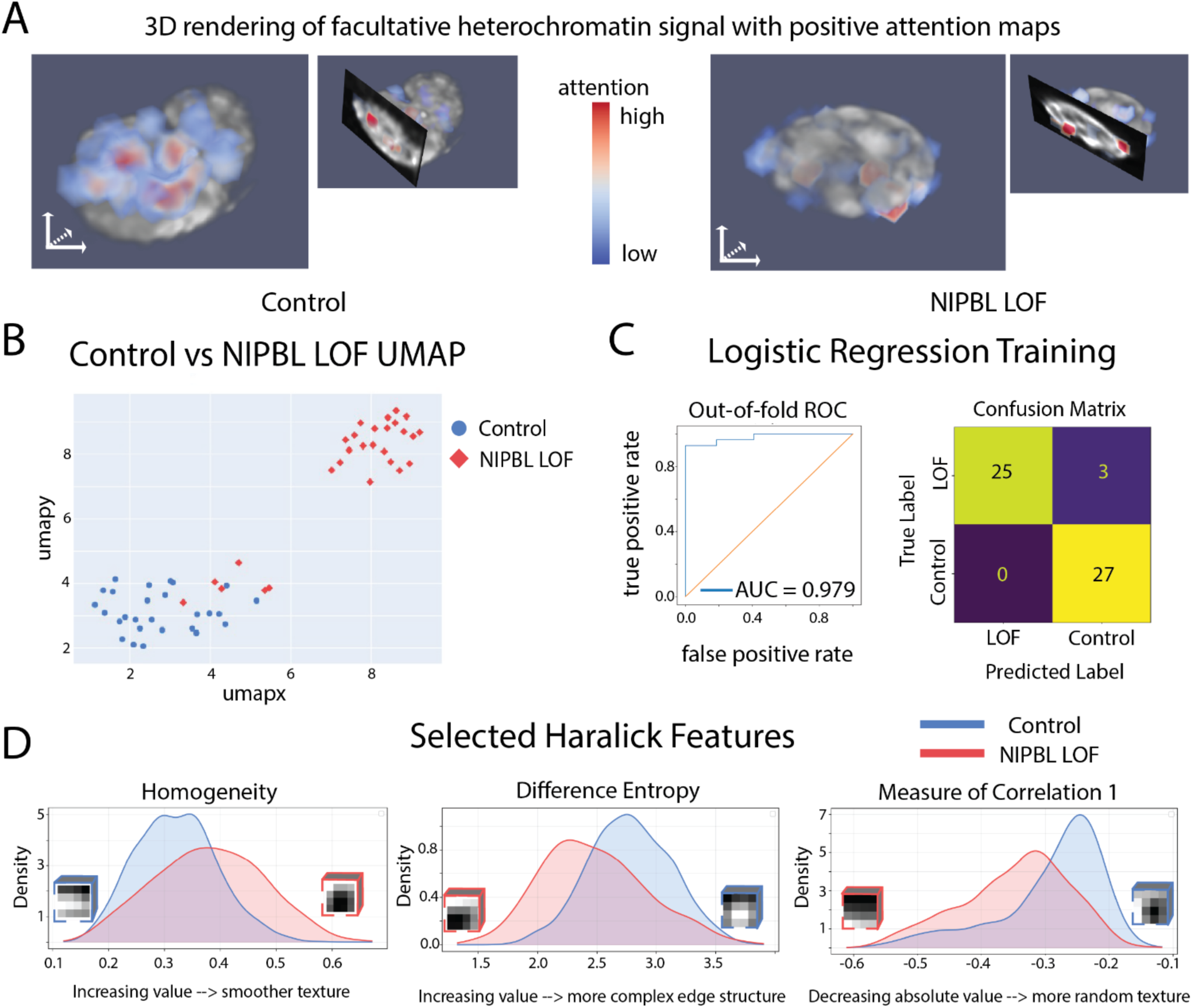
Facultative Heterochromatin biosensor results. A: 3D renderings of the biosensor channel overlaid with the positive attention maps to highlight areas of high attention. B: UMAP results from the normalized image embedding vectors C: A logistic regression model was trained on the image labels and normalized embedding vectors. The model trained with an AUC-ROC of 0.S7S. D: Selected kernel density estimate graphs from various Haralick Features. The KDEs were generated by analyzing patch-level Haralick feature values and summing over the two conditions.

This clustering, the separation between control and loss of function seen in the UMAP, indicated that the algorithm had captured biologically relevant intensity patterns—i.e., some combination of visual words was sufficient to discriminate between control and LOF nuclei. However, the representation itself does not, *a priori*, reveal which features drive the separation. Although one can inspect normalized frequency vectors to identify words that are more prevalent in one condition than the other, the soft assignment (linear combinations of many words per patch) makes it difficult to intuitively understand what common words “look like” in terms of raw image structure. In principle, two patches might differ because one has a high coefficient for word 1 but low for word 2, whereas another has high values for both words, and so on across many words.

To understand which words carried the most discriminative information, we trained a logistic regression (LR) model on the normalized frequency vectors using the known condition labels (control vs. NIPBL LOF). The LR model achieved an area under the receiver-operating characteristic curve (AUC-ROC) of 0.979, confirming that the frequency vectors alone were sufficient for accurate classification (Figure 3C). The LR model also provided a weight (coefficient) for each visual word, quantifying the extent to which that word contributed to predicting the LOF versus control class. Words that predicted the correct class were assigned positive weights, words that predicted the incorrect class were assigned negative weights.

We next asked which regions within each nucleus contributed most strongly to this classification. Using the logistic regression coefficients as weights for the visual words, we computed an “attention” score for each patch and mapped these scores back into the 3D volume to generate attention maps (Figure 3A). To describe these high-attention regions quantitatively, we first thresholded the attention maps to define “high-attention” voxels and performed 3D particle analysis. These maps revealed distinct spatial patterns in control versus LOF nuclei. Control nuclei tended to contain a small number of relatively large, contiguous high-attention regions that spanned substantial portions of the nuclear volume. In contrast, LOF nuclei displayed multiple smaller, more fragmented high-attention regions that were spatially restricted, indicating a shift from broad, extended domains to more fragmented patterns of informative signal (SI Figure 6).

Finally, we examined the local intensity patterns within these high-attention regions using 3D Haralick texture features. Each high-attention patch was summarized by the standard set of 13 Haralick features derived from gray-level co-occurrence statistics. Comparing feature distributions between conditions revealed consistent trends: LOF nuclei exhibited high-attention regions with more smoothly varying textures, greater homogeneity, increased structural regularity, and stronger statistical coupling of neighboring intensities (Figure 3D). Collectively, these metrics indicate that, in the regions most informative for classification, LOF nuclei display smoother, less punctate chromatin-associated signal than control nuclei. Thus, the 3D BoVW framework, coupled with a simple linear classifier and texture analysis, not only separates control and NIPBL LOF nuclei but also points to specific, biologically interpretable differences in chromatin texture underlying this separation.

The primary analysis above focused on the facultative heterochromatin probe, which is enriched in facultative heterochromatin, but the same pipeline was applied independently to the other chromatin markers in the dataset. Hoechst—a commonly used DNA dye that binds the minor groove with preference for AT-rich regions—produces the familiar pattern of bright, compact heterochromatin and dimmer euchromatin. In this dataset, it was the least informative channel: UMAP embeddings of the BoVW frequency vectors showed no separation between control and LOF cells, and the logistic regression model performed poorly (AUC–ROC = 0.716, SI Figure 1).

CTCF labeling, which marks genomic sites involved in higher-order chromatin organization and is enriched at loop anchors and topologically associating domain (TAD) boundaries, showed only weak structure in the embedding. UMAP again revealed minimal separation, although classification improved modestly relative to Hoechst (AUC–ROC = 0.817), suggesting subtle but detectable condition-dependent differences (SI Figure 2).

Finally, H3.3—associated with transcriptionally active chromatin—carried a strong condition-sensitive signal. This channel produced the second-best separation in UMAP (after the facultative heterochromatin probe) and supported robust classifier performance (AUC–ROC = 0.950), consistent with substantial reorganization of features linked to active chromatin under the perturbation (SI Figure 3).

### 5.3 3D Timelapse Results

The receptor clustering timelapse dataset provides a stringent test case because it departs from the “clean” single-cell setting used for the nuclear volumes. Each 3D frame corresponds to an entire field of view containing densely packed neurons with substantial overlap. In addition, confocal acquisition produces pronounced axial anisotropy, and each field of view contains unavoidable biological heterogeneity (cells at different differentiation states and variable expression levels).

Despite these constraints, prior conventional analyses based on manual/algorithmic segmentation of receptor clusters followed by size and morphology measurements showed that ligand addition (netrin) increases the number of guidance receptor Deleted in Colorectal Cancer (DCC) clusters and Partitioning defective 3 (Pard3) overexpression (OE) promotes larger and more persistent membrane-associated DCC clusters^5^. These established effects make the dataset a useful benchmark for representation learning: the distinction between pre- and post-netrin frames is visually apparent and serves as an internal positive control, whereas differences between control and Pard3-overexpressing samples are subtler and provide a more stringent test of sensitivity to perturbation-induced phenotypes. Because the dataset has been previously analyzed with handcrafted features and segmentation, we could also evaluate whether the 3D BoVW representation recapitulates known results without explicit segmentation.

As expected, the most prominent separation identified by the algorithm, visualized with UMAP in Figure 4B, was between pre- and post-netrin images. Beyond this, the method also distinguished control cells from Pard3-overexpressing cells, indicating that SIFT keypoints encoded as visual words captured sufficient information to replicate, in an unsupervised manner, results that previously required manual segmentation and feature extraction.

**Figure 4.**
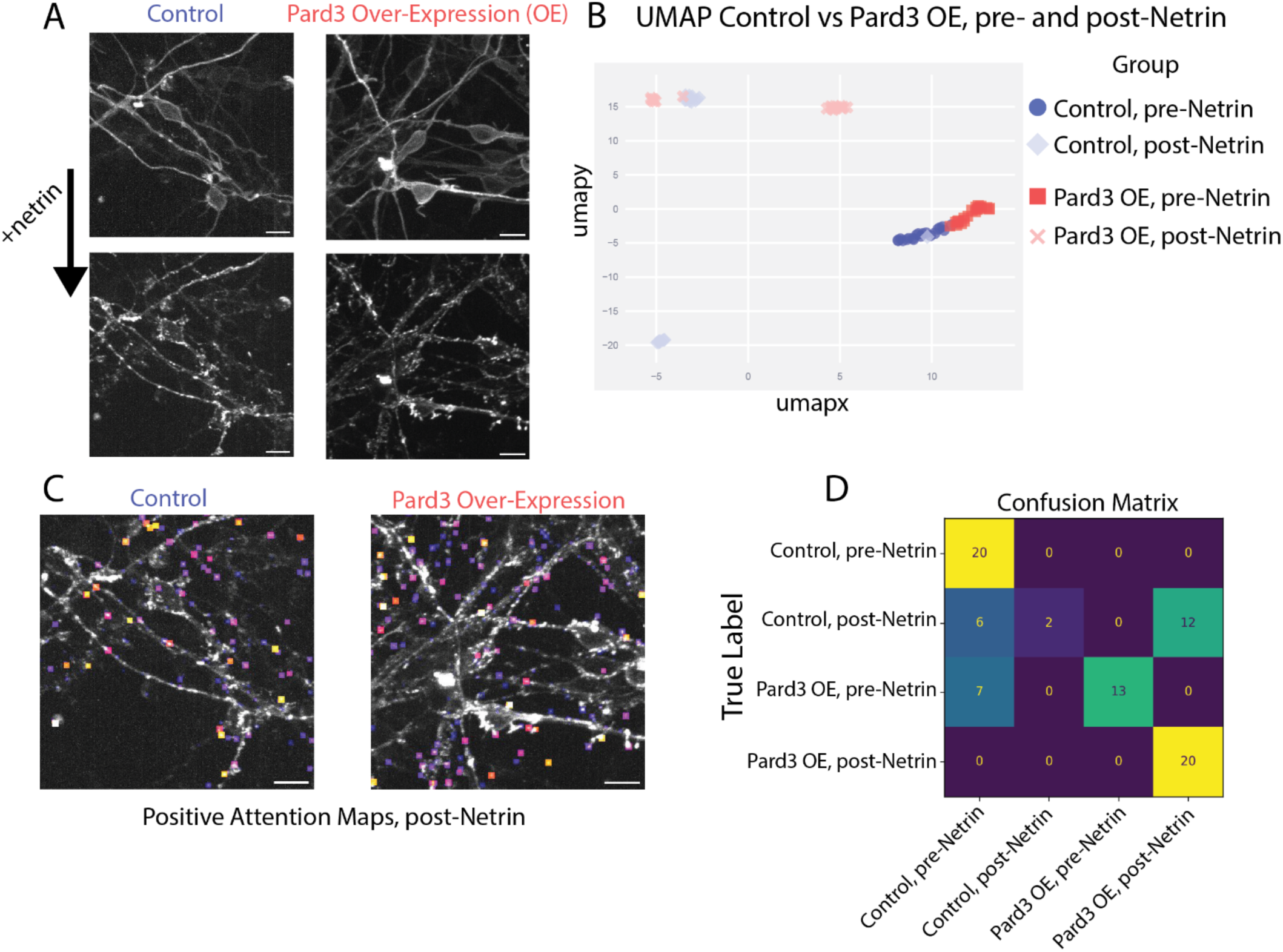
DCC Timelapse dataset results. A: representative images from the data showing CGN cultured with and without Pard3 overexpression [scale bar = 10 µm] before and after addition of the ligand netrin. B: UMAP graph of image embedding vectors. C: Representative images overlaid with the positive attention maps. D: Confusion matrix after training a 4-class logistic regression model.

The clustering results also reflected underlying biological complexity. The overexpression of Pard3 was achieved by the nucleofection of the Pard3 plasmid. As a result, although all Pard3-overexpressing cells expressed more Pard3 than controls, the magnitude of overexpression varied, producing overlap between conditions in both pre-and post-netrin images.

Temporal heterogeneity was likewise evident in the low-dimensional embedding (UMAP). Post-netrin data points that clustered near pre-netrin points corresponded to the earliest time frames after netrin addition. Because timelapse imaging involved multiple fields of view, some cells were exposed to netrin slightly longer than others before the first post-stimulation frame was acquired, leading to a continuum of response states rather than a sharp pre/post boundary.

Feeding the normalized histograms into a four-class logistic regression model revealed which conditions were easier to classify and which conditions were more confusing. The model made no errors when classifying the control pre-netrin images and the Pard3 OE post-netrin images but had more difficulty with the other two classes (Figure 4D).

Together, these results show that the 3D BoVW framework *(i)* captured the expected large-scale ligand-driven change (pre-vs post-netrin), *(ii)* resolved more subtle differences associated with polarity pathway perturbation (control vs Pard3 overexpression), and *(iii)* encoded both biological heterogeneity (variable expression) and temporal heterogeneity (variable time after stimulation). A remaining limitation is that analysis is performed at the level of whole fields of view rather than individual cells; given dense cultures and mixed developmental states, field-level aggregation likely contributes to the partial overlap between conditions and sets an upper bound on separability in this dataset.

## 6. Discussion

The design of this pipeline was guided by the goal of creating a broadly applicable tool for analyzing complex microscopy datasets, rather than a method tailored exclusively to one specific dataset. By operating directly on 3D image structure and learning data-driven visual vocabularies, the framework is well suited to problems in which differences between conditions are visually apparent to an expert but resist simple, hand-crafted quantification.

One natural application is screening. In experiments with many genetic perturbations, the pipeline can be used to generate image-level embeddings and quantify how strongly each condition deviates from a control distribution. Ranking conditions by their divergence from control can help prioritize perturbations for follow-up studies. A similar strategy could be applied across imaging channels, for example to identify which marker exhibits the most discriminative phenotype in a multi-channel experiment.

Beyond screening, the method can assist in interpreting which aspects of image structure distinguish conditions. Because the downstream classifier operates on interpretable Bag-of-Visual-Words features, its weights can be mapped back to image regions and intensity textures that drive classification. In this sense, the algorithm is intended to complement, not replace, expert evaluation: it provides quantitative summaries and spatial attributions that can sharpen and guide biological interpretation.

The approach also has limitations. First, feature extraction is driven by SIFT-like keypoints in the intensity landscape, making the method inherently intensity-based. In datasets where the signal is extremely homogeneous, with few local maxima or minima, keypoint detection—and therefore the BoVW representation—may fail to capture meaningful variation. Second, in its current form the pipeline operates on a single channel at a time. It does not directly model relationships between channels, and therefore cannot exploit information that is present only in the joint spatial organization or colocalization of multiple markers.

Despite these constraints, applying the pipeline independently to multiple chromatin markers proved informative, because each channel emphasized a different aspect of nuclear organization. Hoechst provided little discriminative signal: UMAP showed minimal separation and the logistic regression model performed poorly, suggesting that overall DNA distribution (heterochromatin vs euchromatin) was largely unchanged between conditions (or that any changes were not well captured by an intensity-keypoint representation). In contrast, H3.3—associated with transcriptionally active chromatin—contained a richer and more condition-sensitive structure, yielding clear separation in UMAP and supporting classifier training. The primary probe, for facultative heterochromatin, similarly carried strong discriminatory information, producing robust separation and model performance, whereas CTCF fell in between, with weaker but detectable separation and modest classification accuracy.

Viewed biologically, this channel-by-channel behavior is itself a useful readout. NIPBL knockdown is expected to disrupt cohesin-mediated loop extrusion and thereby alter gene regulation and chromatin architecture. The pronounced changes observed in H3.3 are consistent with a strong impact on transcriptionally active regions. Interestingly, facultative heterochromatin also showed marked differences, even though it is enriched in facultative heterochromatin. This suggests that the perturbation affects nuclear architecture broadly, producing measurable reorganization even in regions not classically considered transcriptionally active.

More generally, running the same analysis across multiple markers transforms a “single-score” classification into a comparative, mechanistic probe: channels that separate well likely report on aspects of organization most affected by the perturbation, while channels that do not may indicate relative stability (or limited sensitivity of the representation for that signal type). This multi-channel perspective therefore increases the biological insight gained from the same set of nuclei, enabling a more nuanced interpretation of how different chromatin compartments respond to genetic perturbation.

The chromatin architecture dataset presented here illustrates an ideal use case. An expert observer can readily see that NIPBL knockdown alters nuclear morphology and chromatin organization, but an *a priori* feature set to quantify these differences is not obvious. The pipeline identifies image regions and texture patterns that are most informative for distinguishing control and knockdown nuclei. In NIPBL-depleted cells, cohesin dysfunction produces enlarged nuclei with large chromatin-poor “voids,” effectively compressing chromatin into a smaller accessible volume. The learned features emphasize smoother, more homogeneous chromatin signal within these restricted regions, consistent with a model in which chromatin is packed more uniformly into the remaining territory. In contrast, control nuclei exhibit a more punctate and heterogeneous chromatin texture distributed over a larger volume, which the classifier associates with the control condition. Thus, the quantitative output of the algorithm aligns with and refines the qualitative biological interpretation.

Across datasets, the pipeline was built following a “minimal complexity” principle: we began with the simplest variant capable of separating conditions and introduced additional components only as needed. For the timelapse DCC clustering dataset, SIFT keypoint detection combined with a 3D BoVW representation was sufficient to distinguish not only the presence or absence of ligand (netrin) but also the more subtle effect of Pard3 overexpression on receptor organization.

To assess whether a more complex deep-learning representation offered an advantage, we trained a masked autoencoder (MAE) on the same nuclear dataset and used the learned embeddings for downstream classification. The MAE-based embeddings underperformed our 3D BoVW representation (SI Fig. 8; AUC-ROC 0.837 for 3D MAE and 0.666 for 2D projection MAE, versus 0.979 for the facultative heterochromatin BoVW analysis) reinforcing our choice of a simpler pipeline that not only performed better in this setting but also retained direct interpretability through mappable visual words and attention maps.

All code for the 3D pipeline is made publicly available on the 3D BoVW repository on GitHub [https://github.com/PittmanAEP/3DBoVW], enabling researchers to apply, adapt, and extend these tools to diverse imaging modalities and biological questions.

## Acknowledgements

We are indebted to Sharon King and Rebecca Petersen of the Department of Developmental Neurobiology Neuroimaging Laboratory at St. Jude Children’s Research Hospital for maintaining and aligning the instruments used in this study’s lattice light-sheet imaging sessions. The Solecki lab is funded by the American Lebanese Syrian Associated Charities and by grants R01 NS066936, R01 NS104029, and R01 NS139519 from the National Institute of Neurological Disorders. The content of this manuscript is solely the responsibility of the authors and does not necessarily represent the official views of the NIH.

## SI Figures

**SI Figure 1.**
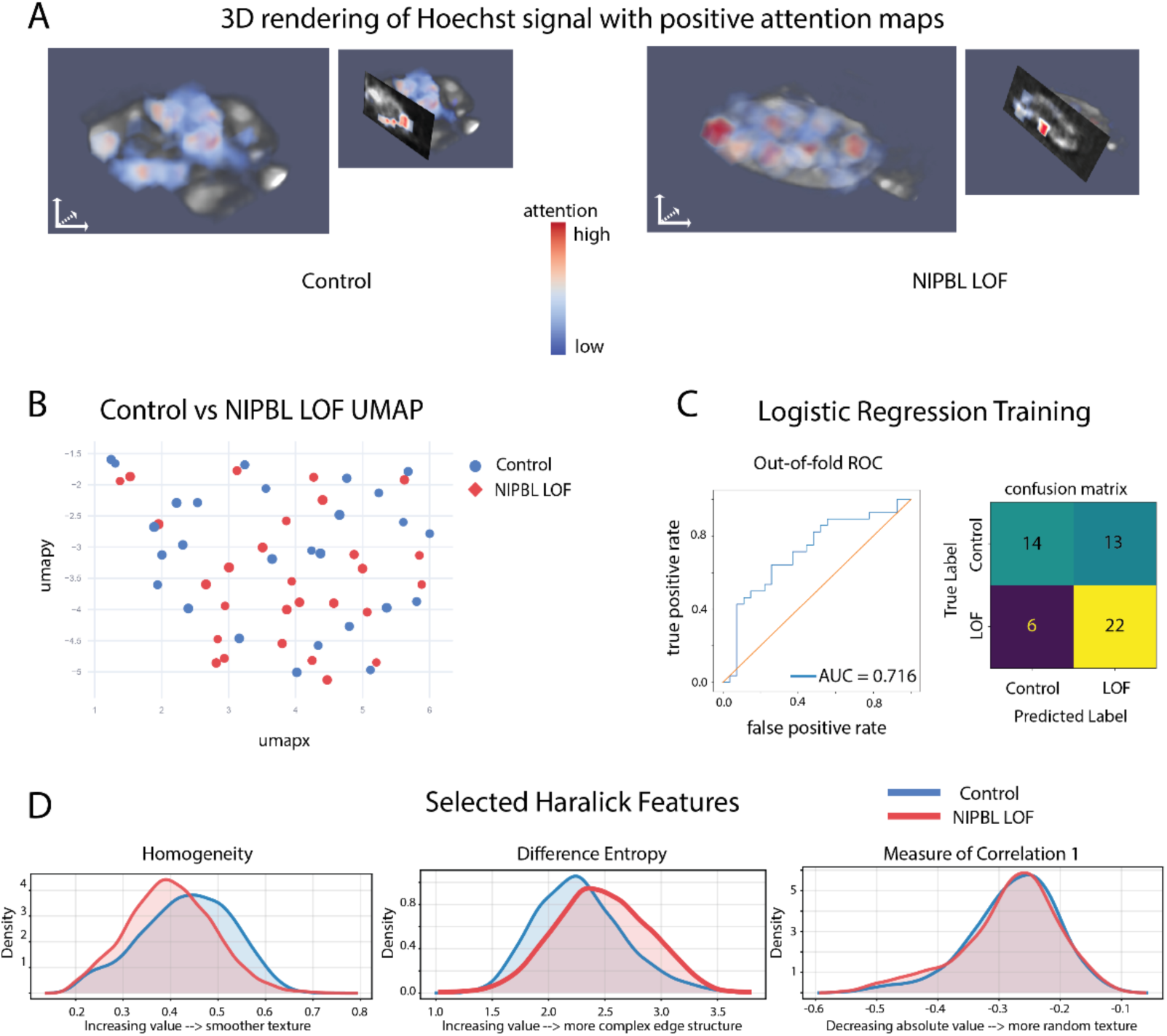
Hoechst results. A: 3D renderings of the Hoechst channel overlaid with the positive attention maps to highlight areas of high attention. B: UMAP results from the normalized image embedding vectors C: A logistic regression model was trained on the image labels and normalized embedding vectors. The model trained with an AUC-ROC of 0.716. D: Selected kernel density estimate graphs from various Haralick Features. The KDEs were generated by analyzing patch-level Haralick feature values and summing over the two conditions.

**SI Figure 2.**
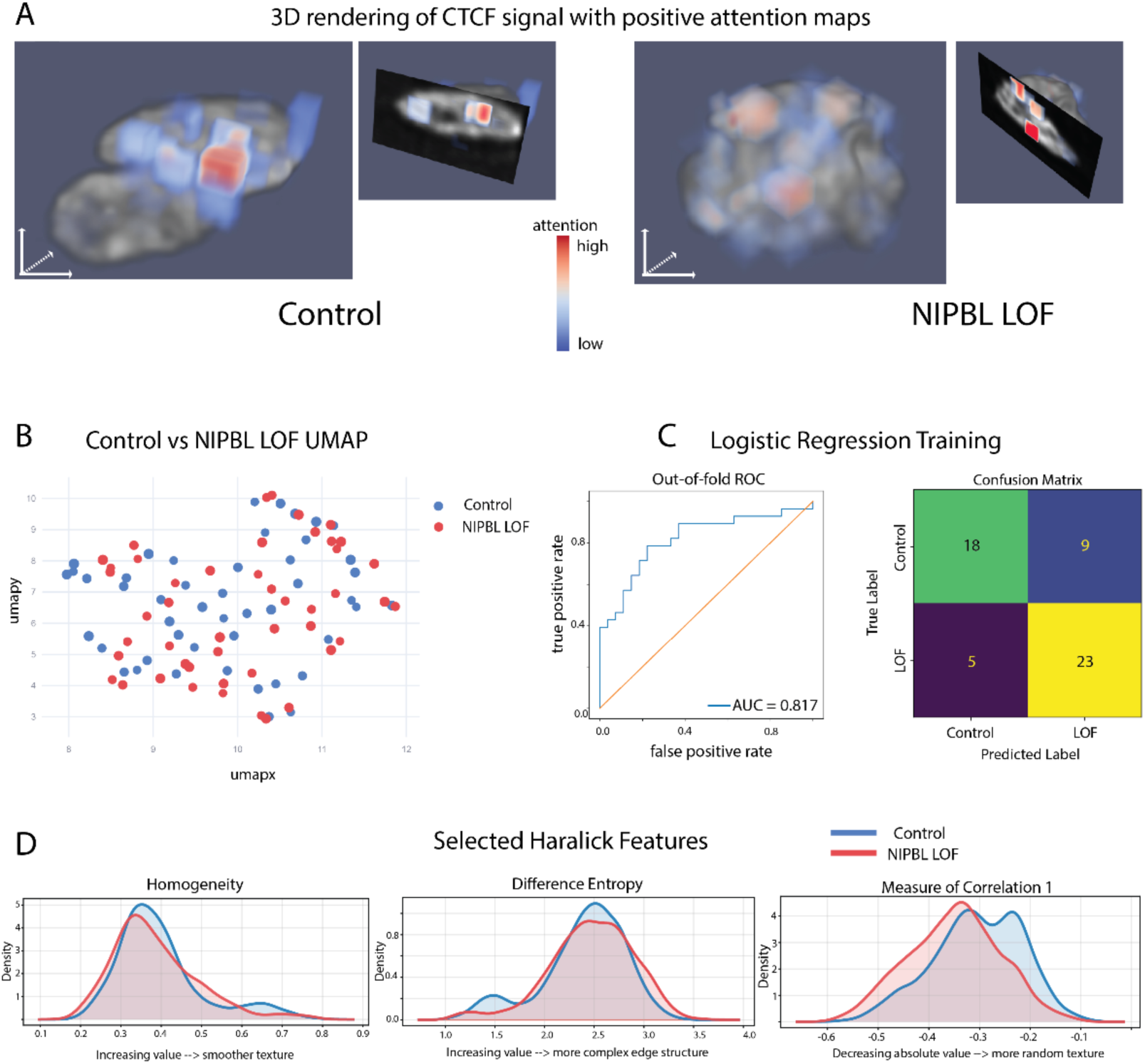
CTCF Results. A: 3D renderings of the CTCF channel overlaid with the positive attention maps to highlight areas of high attention. B: UMAP results from the normalized image embedding vectors C: A logistic regression model was trained on the image labels and normalized embedding vectors. The model trained with an AUC-ROC of 0.817. D: Selected kernel density estimate graphs from various Haralick Features. The KDEs were generated by analyzing patch-level Haralick feature values and summing over the two conditions.

**SI Figure 3.**
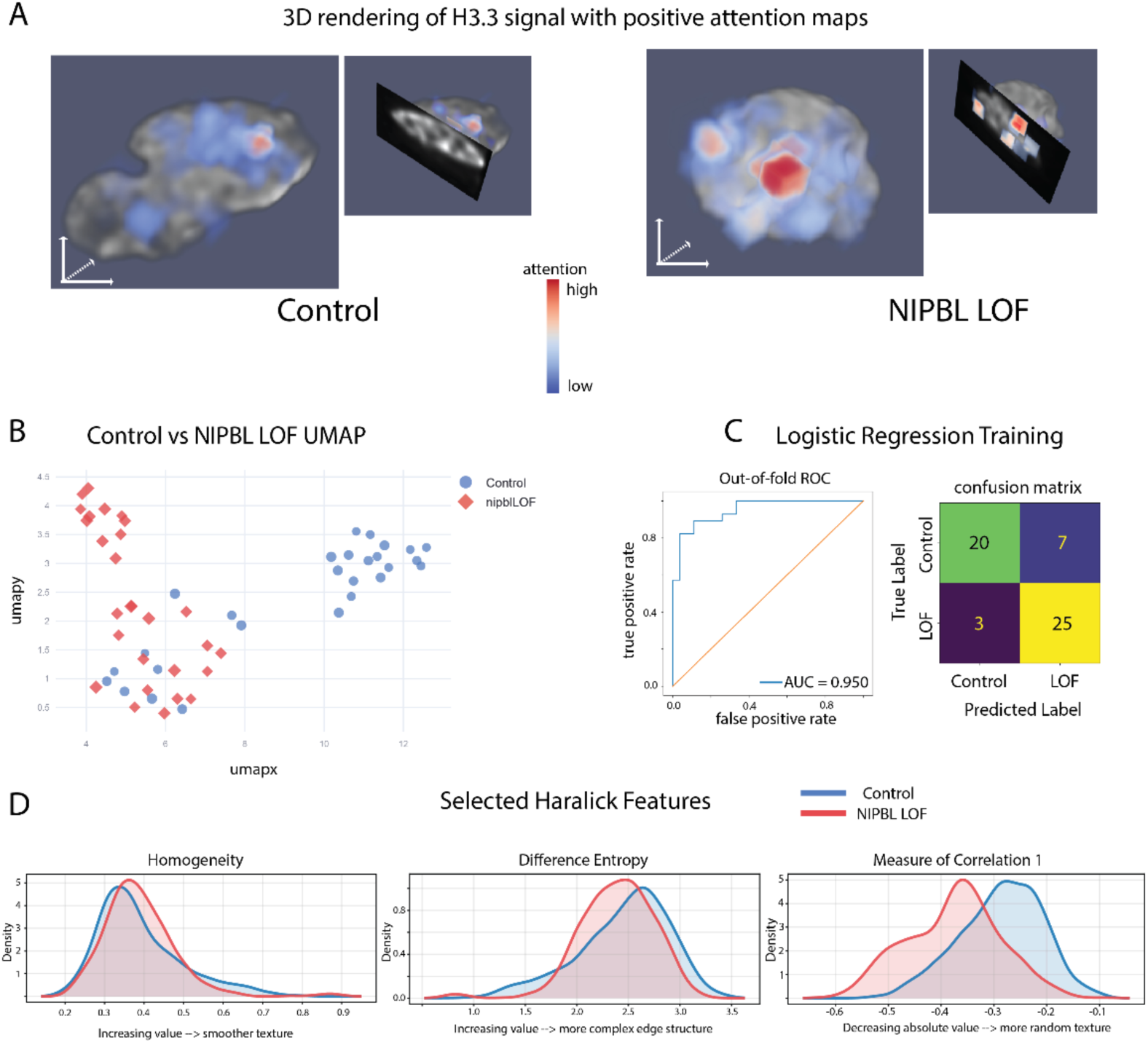
H3.3 Results. A: 3D renderings of the H3.3 channel overlaid with the positive attention maps to highlight areas of high attention. B: UMAP results from the normalized image embedding vectors C: A logistic regression model was trained on the image labels and normalized embedding vectors. The model trained with an AUC-ROC of 0.S50. D: Selected kernel density estimate graphs from various Haralick Features. The KDEs were generated by analyzing patch-level Haralick feature values and summing over the two conditions.

**SI Figure 4.**
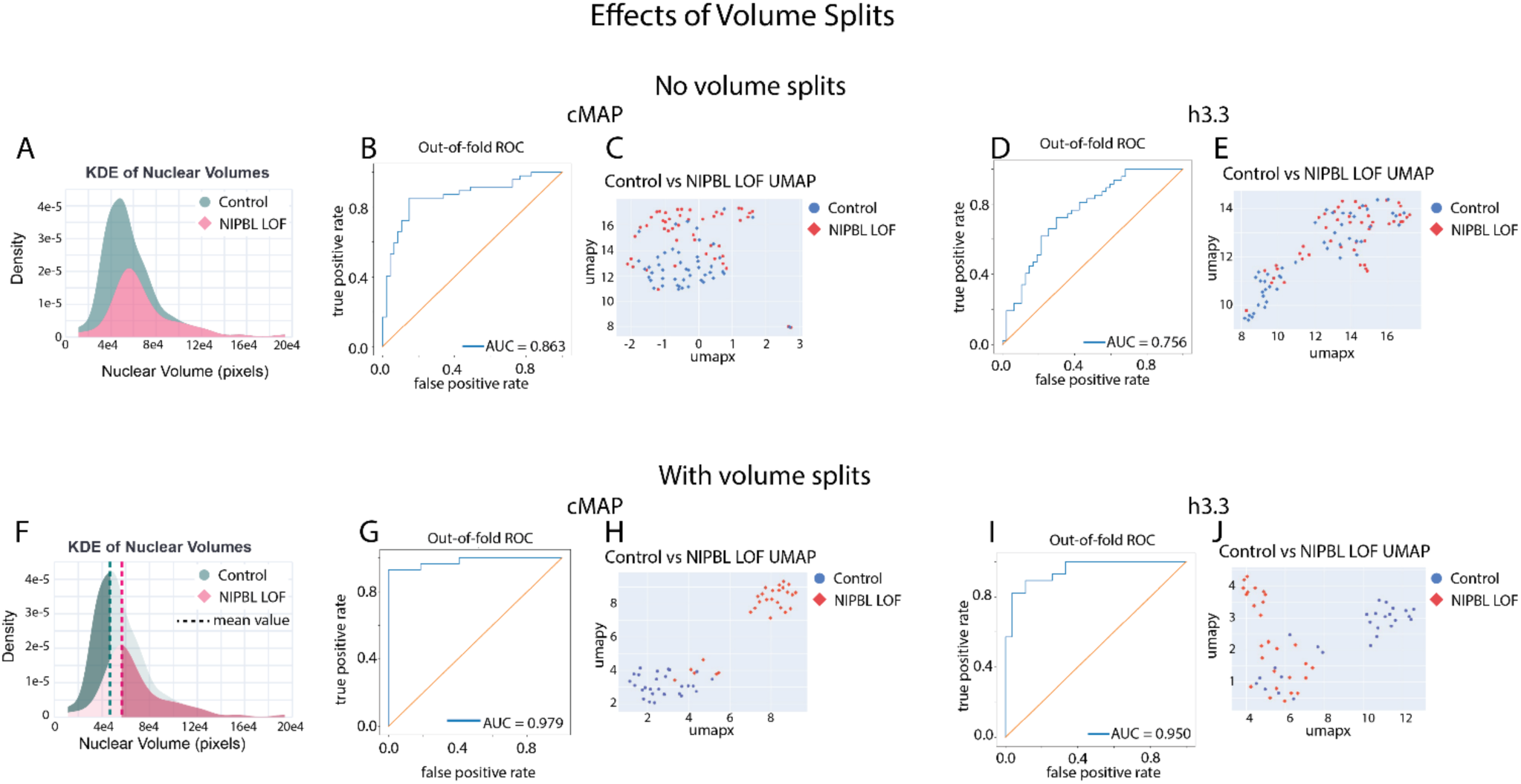
Effects of volume splits on classification. Analyses were performed either on the full dataset without volume splitting (top row, A–E) or after size stratification (bottom row, F–J), using smaller control nuclei and larger NIPBL LOF nuclei split at the median nuclear volume, as described for the chromatin dataset analyses. A and F, kernel density estimates of nuclear volume distributions for control and LOF nuclei; F shows the mean volume for each condition and the applied split. B and G, out-of-fold ROC curves for logistic regression models trained on normalized BoVW vectors from the facultative heterochromatin channel, with AUC increasing from 0.8C3 to 0.S7S after stratification. C and H, corresponding UMAP embeddings for facultative heterochromatin. D and I, out-of-fold ROC curves for logistic regression models trained on H3.3 BoVW vectors, with AUC increasing from 0.75C to 0.S50 after stratification. E and J, corresponding UMAP embeddings for H3.3. These results show that volume-based stratification reduces overlap between control and NIPBL LOF nuclei and improves condition separation in both channels.

**SI Figure 5.**
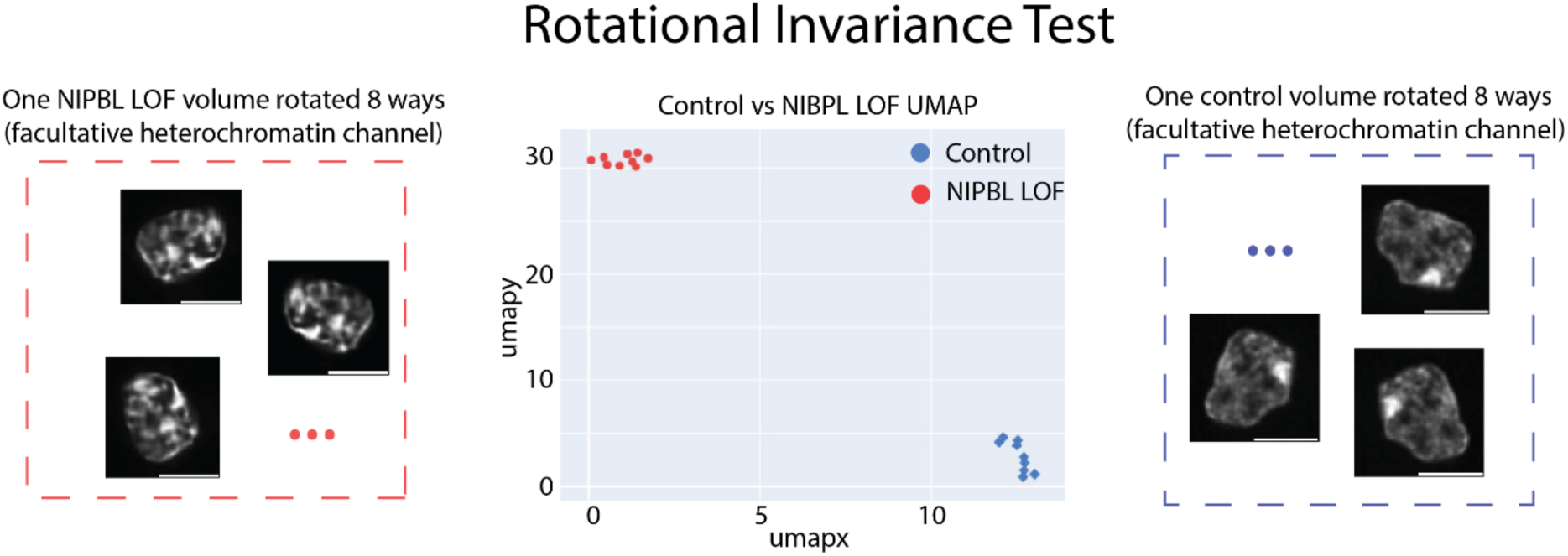
Rotational Invariance. A minimal test dataset was generated from two individual facultative heterochromatin biosensor nucleus crops, one control and one NIPBL LOF. For each volume, eight rotated variants were created and analyzed together with the original image using the full pipeline. The resulting UMAP embedding shows that rotated versions cluster with their corresponding source image rather than separating by orientation, consistent with the rotationally robust descriptor design described in the manuscript

**SI Figure 6.**
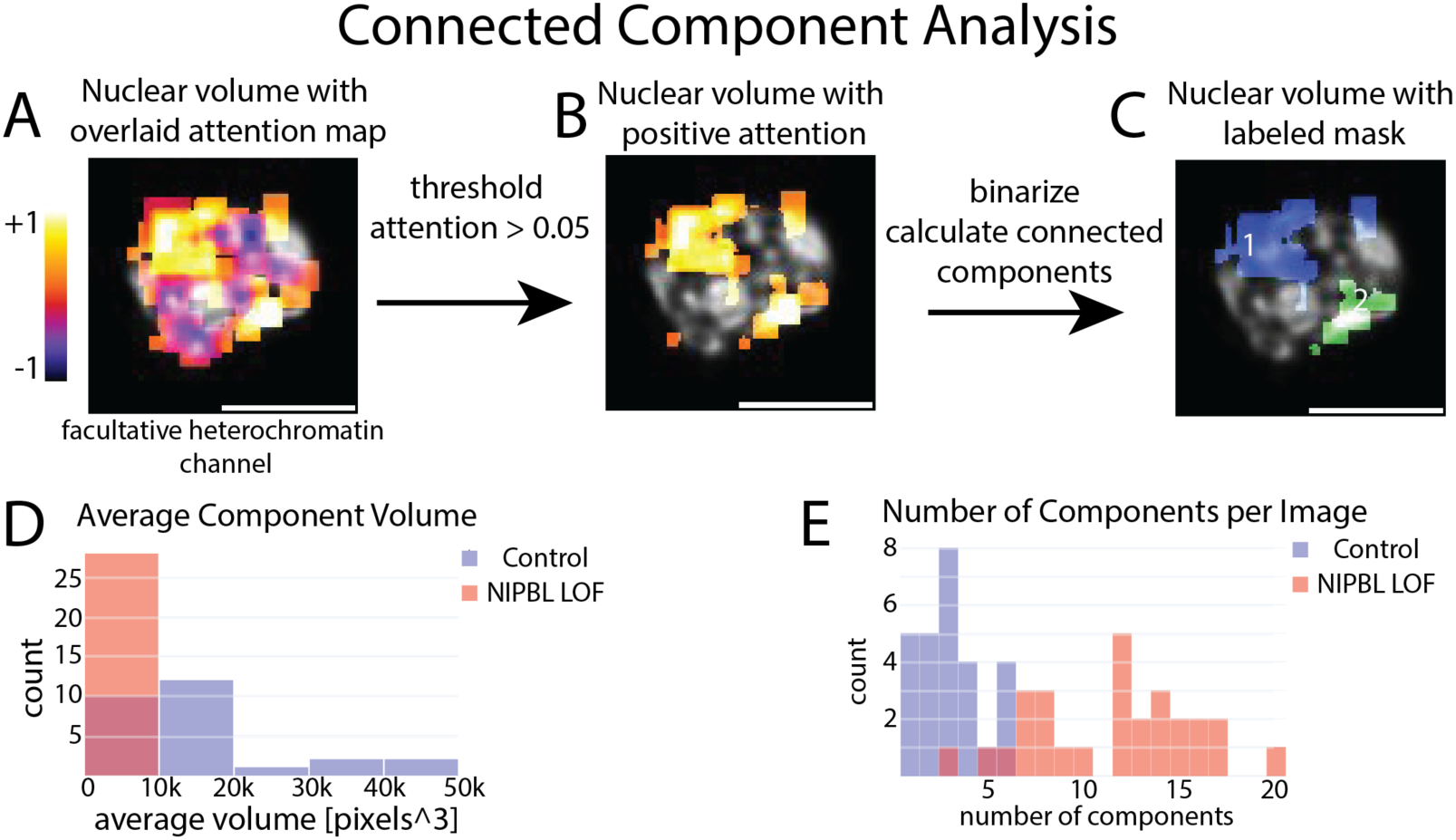
Connected-component (“blob”) analysis of positive attention regions in facultative heterochromatin biosensor expressing CGNs. A, representative nucleus overlaid with the attention map, with positive and negative patch contributions shown on a signed scale. B, positive attention regions retained after thresholding the attention map (> 0.05). C, binary mask after connected-component labeling, with individual high-attention components identified for downstream quantification. D, distribution of average connected-component volume per image for control and NIPBL LOF nuclei. E, distribution of the number of connected components per image for each condition. Consistent with the attention-map trends described in the manuscript, control nuclei tended to contain fewer, larger connected high-attention regions, whereas NIPBL LOF nuclei showed a greater number of smaller, more fragmented high-attention regions

**SI Figure 7.**
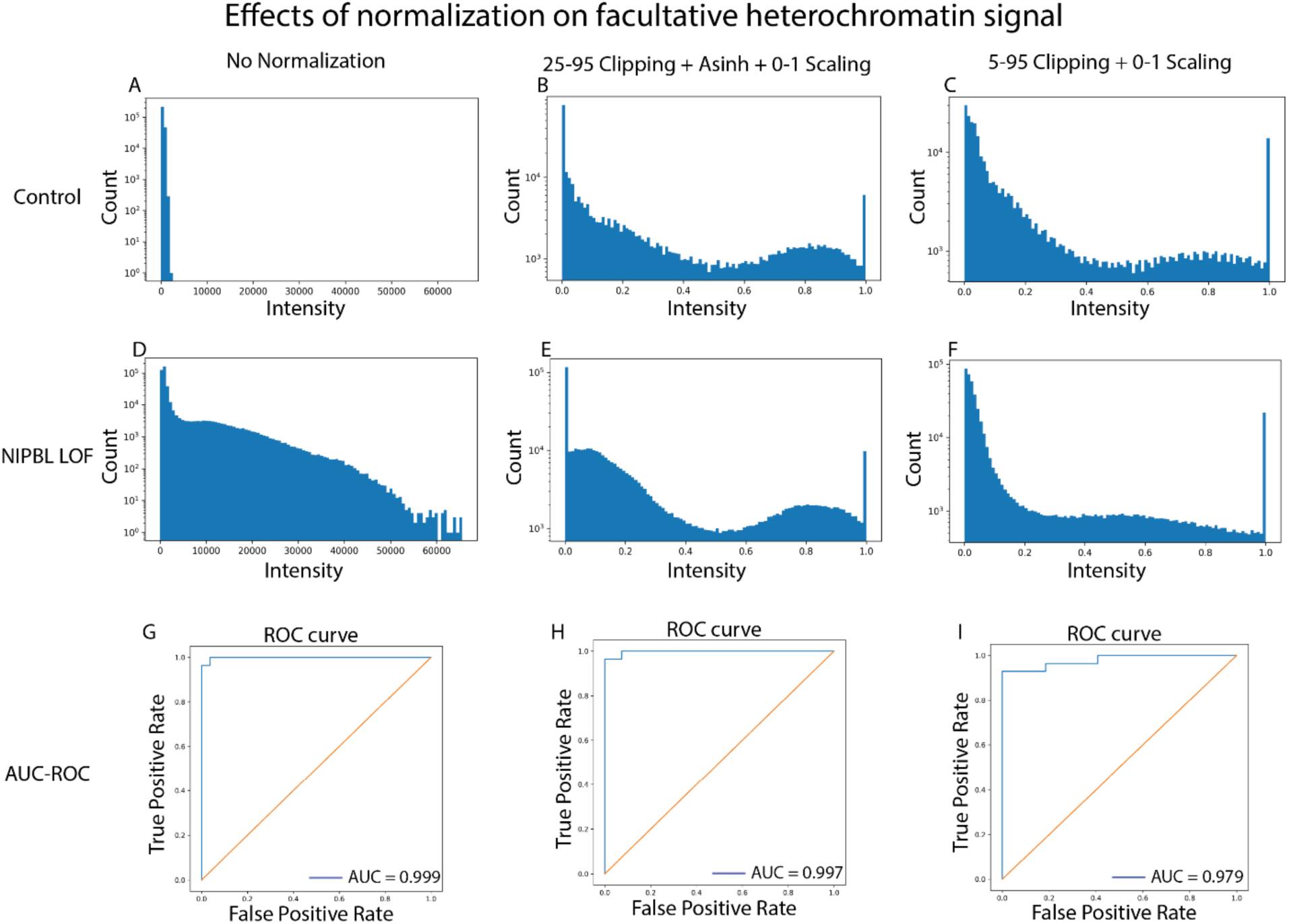
Effects of normalization on image intensity. A-C, intensity histograms of the facultative heterochromatin channel from a selected control image under various normalization conditions. D-F, intensity histograms of the facultative heterochromatin channel from a selected NIPBL LOF image under various normalization conditions. G-I, ROC curves from training the LR model on the entire dataset under various normalization conditions.

**SI Figure 8.**
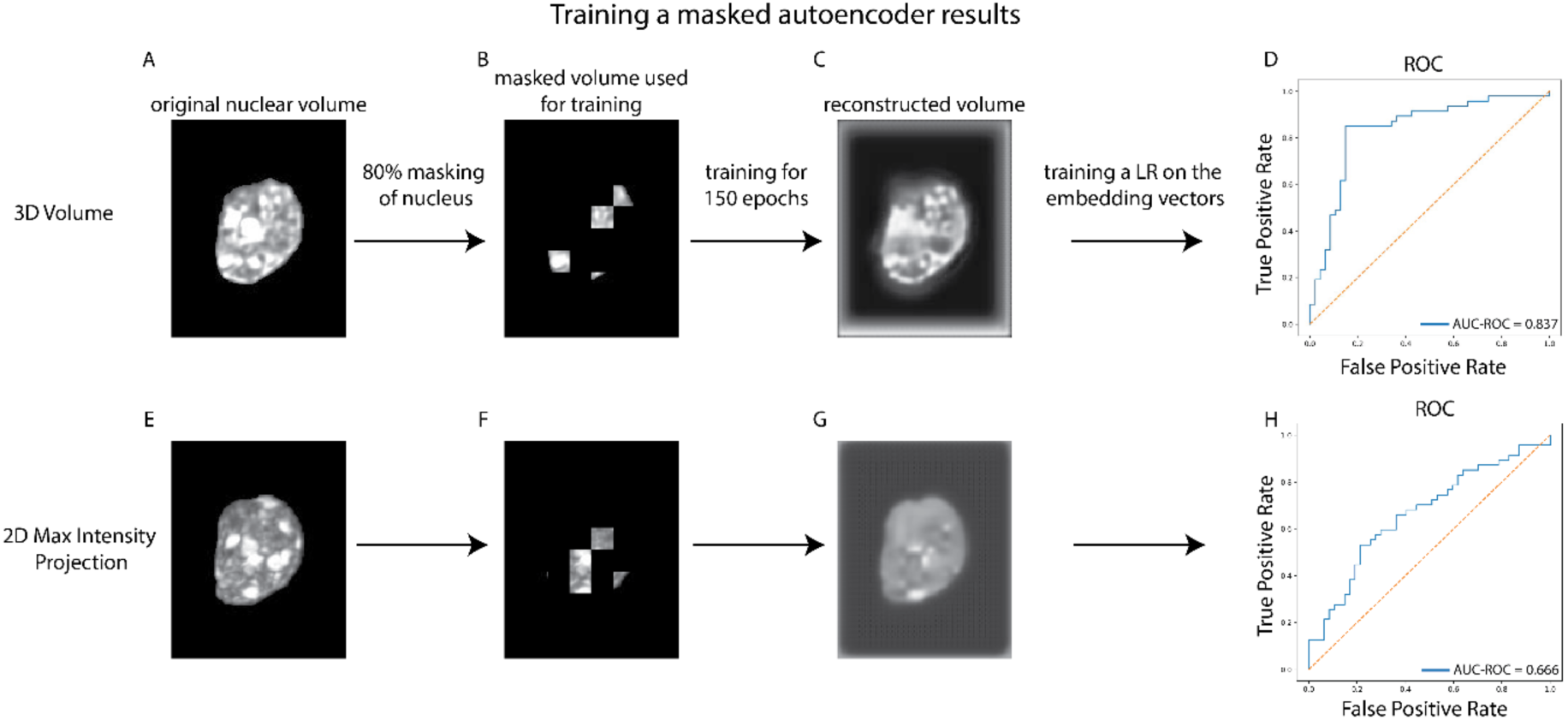
Comparison with embedding vectors from custom trained masked autoencoder. Full facultative heterochromatin biosensor channel nuclear volumes, or 2D maximum intensity projections, were used to train a custom masked autoencoder (MAE) to generate embedding vectors to compare to the embedding vectors from the Bag-of-visual-words pipeline. The MAE was implemented as a 3D convolutional masked autoencoder (ConvMAE3D; base width 64) trained with 80% blockwise masking (4 × 1C × 1C voxels) for 150 epochs using AdamW (learning rate = 1 × 10⁻⁴) and masked L1 reconstruction loss; image-level embeddings were obtained by mean-pooling bottleneck features for downstream logistic regression. A,C: original images (volumetric or 2D max projection). B,F: images after 80% masking was done. Only the nucleus was masked to avoid having the MAE learn background textures. C,G: reconstruction results after 150 epochs of training. D,H: the embedding vectors from the trained MAE were used to train a 2-class logistic regression model in the same way as with the BoVW vectors in the paper. Both cases (volumetric and 2D) did not perform as well as the BoVW vectors.

